# scRNA-seq reveals persistent aberrant differentiation of nasal epithelium driven by TNFα and TGFβ in post-COVID syndrome

**DOI:** 10.1101/2024.01.10.574801

**Authors:** A. Fähnrich, K.D. Reddy, F. Ott, Y. Maluje, R. Saurabh, A. Schaaf, S. Winkelmann, B. Voß, M. Laudien, T. Bahmer, Jan Heyckendorf, F. Brinkmann, S. Schreiber, W. Lieb, M. Weckmann, H. Busch

## Abstract

Post-COVID syndrome (PCS) currently affects approximately 3-17% of people following severe acute respiratory syndrome coronavirus 2 (SARS-CoV-2) infection and has the potential to become a significant global health burden. PCS presents with various symptoms, and methods for improved PCS assessment are presently developed to guide therapy. Nevertheless, there are few mechanistic insights and treatment options. Here, we performed single-cell RNA transcriptomics on nasal biopsies from 33 patients suffering from PCS with mild, moderate, or severe symptoms. We identified 17 different cell clusters representing 12 unique cell populations, including all major epithelial cell types of the conducting airways and basal, secretory, and ciliated cells. Severe PCS was associated with decreased numbers of ciliated cells and the presence of immune cells. Ensuing inflammatory signaling upregulated TGFβ and induced an epithelial-mesenchymal transition, which led to the high abundance of basal cells and a mis-stratified epithelium. We confirmed the results *in vitro* using an air-liquid interface culture and validated TNFα as the causal inflammatory cytokine. In summary, our results show that one mechanism for sustained PCS is not through continued viral load, but through the presence of immune cells in nasal tissue leading to impaired mucosal barrier function and repeated infections. These findings could be further explored as a therapeutic option akin to other chronic inflammatory diseases by inhibiting the TNFα-TGFβ axis, restoring the nasal epithelium, and reducing respiratory tract-related infections.

## Introduction

Severe acute respiratory syndrome coronavirus 2 (SARS-CoV-2) is the viral agent that causes COVID-19 (coronavirus disease 2019) and was discovered in China in December 2019. By now, the pathophysiology and molecular mechanisms have been studied in detail using epidemiology, multi-Omics approaches, single-cell and *in vitro* models [1, 2]. The viral tropism of SARS-CoV-2 is determined by the abundance and co-expression of two key cell-surface proteases, such as angiotensin-converting enzyme 2 (ACE2), transmembrane serine protease 2 (TMPRSS2). These proteins are found highly expressed in the airway epithelium, which forms the first barrier against environmental exposures [3-5]. As such, the multi-ciliated cells of the upper respiratory tract likely present the first site of SARS-CoV-2 infection. However, as viral load increases the lower respiratory tract also presents as a key entry site, due to high expression of ACE2 and TMPRSS2 on alveolar type 2 (AT2) cells. This causes alveolar damage, inflammation, and an influx of immune cells, driving disease progression and causing worse health outcomes [6]. Simultaneously, a high viral load in the nasal epithelium promotes transmissibility through coughing or sneezing, maximizing SARS-CoV-2’s ability to spread and presenting the nasal epithelium as a primary pathological site [7].

Although most individuals infected with SARS-CoV-2 typically experience low-grade symptoms and recover within a few weeks, a substantial number, estimated in January 2023 at around 65 million globally, continue to cope with long-lasting symptoms [8, 9]. This persistent post-infection multisystem condition, often referred to as “Post-COVID Syndrome” (PCS) or “Long COVID,” is marked by prevalent symptoms such as fatigue, shortness of breath, and cognitive dysfunction. These symptoms significantly impede an individual’s ability to engage in daily activities for extended periods, ranging from months to years [8, 9]. Notably, approximately 10–20% of cases, across all age groups, including children, are estimated to experience PCS. The significance of PCS as a pertinent pathology is validated by the WHO classification as an internationally classified disease (ICD-10 code).

PCS poses a significant threat to individual health, impeding clinical interventions and efforts to improve patient health outcomes. Various hypotheses regarding its cause, such as immune dysregulation, microbiota, neuronal signaling dysfunction, viral reservoirs, and T-cell exhaustion, have been explored in detail by Davis et al [9]. However, gaps exist in our knowledge regarding PCS susceptibility factors, biomarkers, and pathological mechanisms. A deeper understanding of the molecular basis of PCS will advance its diagnosis and treatment.

The nasal epithelium is specialized for respiratory functions, involving air filtration, humidification, and pathogen protection. Basal cells differentiate into various cell types, collectively producing mucus to trap foreign particles and pathogens. Crucially, basal cells continuously regenerate, maintaining a healthy nasal epithelial layer. SARS-CoV-2 primarily infects ciliated epithelial cells and replicates more efficiently in nasal conditions than in the lower respiratory tract [10]. The NAPKON-POP study platform has been initiated in Germany to facilitate population-based PCS studies within the general population and hosts the COVIDOM study [11]. The platform classifies PCS severity based on enduring symptoms that impact quality of life [8]. While this approach based on patient-reported symptoms is beneficial for clinical categorization, associations with certain molecular mechanistic have not been identified so far. Ongoing biomarker research holds promise for the identification of effective markers, potentially enhancing our understanding and treatment strategies for PCS once these markers are identified and comprehensively understood.

As such, to better understand the pathological mechanisms of PCS, we hypothesize a residual disease-state in the nasal epithelia in post-COVID patients. For this, we extended the NAPKON examination protocol and obtained nasal biopsies of n=33 study participants with mild, moderate, and severe PCS. We aimed to identify changes in cellular abundance and interactions that occur post-acute SARS-CoV-2 infection that differ between patients suffering varying clinical presentations of PCS. Single-cell analysis revealed permanent epithelial de-differentiation with a high abundance of basal cells that correlated with symptom severity similar to the state of acute infection despite the absence of a virus load. This is caused by the presence and communication of immune cells and secretion of TGFβ, which impairs proper differentiation of basal cells through induction of epithelial-mesenchymal transition, which we confirmed in a human nasal epithelium model.

## Methodology

### Study population and Sample collection

We obtained nasal biopsies from n=33 patients anterior and medial to the head of the middle turbinate under endoscopic guidance (Figure 1). This standard procedure using a curette scrapes out well-defined biopsies containing epithelial and also cells from lower layers. Patients were registered in the NAPKON-POP cohort and had provided written and informed consent before biopsy collection, aligning with ethical approval. The criteria for patient inclusion were (i) persistence of COVID-19 symptoms for more than three months, (ii) post-acute infection symptom development, and (iii) a worsening of pre-existing comorbidities. In line with a recently established scoring for PCS [8], we categorized study participants into three different post-COVID syndrome groups according to their symptoms: mild (n=5, PCS score < 10.75), moderate (n=11, 10.75 < PCS score < 26.25), and severe (n=17, PCS score > 26.25) [8].

**Figure 1:**
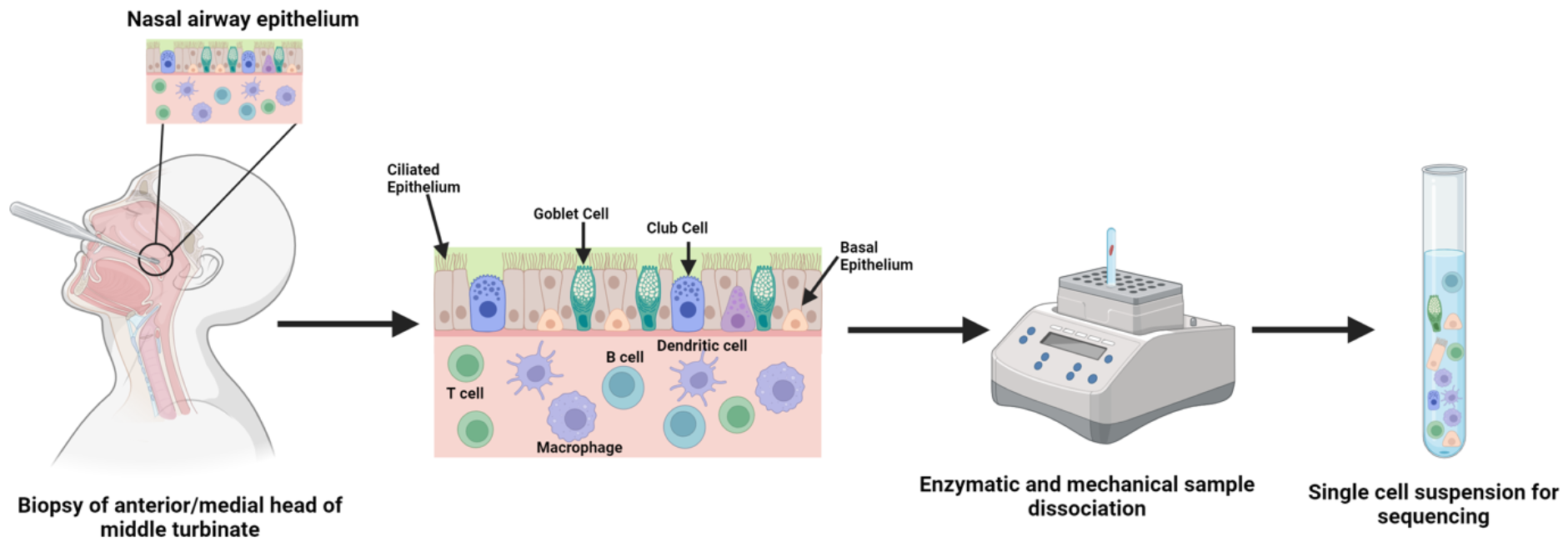
Sample collection and processing. Generated with BioRender.

### Single-cell library prep and sequencing parameters

Single-cell sequencing was performed in collaboration with the Singleron Company, Cologne, Germany, whose workflow allows storage of tissue specimens up to 72 hours before processing and library preparation, ensuring high sample integrity. Nasal curettage samples were collected from study participants at the University Hospital Schleswig-Holstein (UKSH), Campus Kiel, Germany, during routine patient visits. Isolated cell suspensions were shipped overnight to Singleron labs in Cologne, Germany, where the samples were digested into single-cell suspensions using sCelLiVE Tissue Dissociation Solution and library preparation through GEXSCOPE Single Cell RNA Library Kit on a microwell chip (SCOPE-chip) and barcoded beads. Cells were subsequently lysed and the beads with attached poly-A tailed transcripts extracted. The captured mRNA was then reverse transcribed into cDNA, amplified, fragmented, and then ligated with sequencing adapters. The finished library was paired-end sequenced on an Illumina NovaSeq.

### Single-cell RNA-Seq analysis pipeline

Raw gene expression matrices were generated for each sample by a custom pipeline combining *‘kallisto’* (v.0.46.1) and *‘bustools’* (v.0.46.1) using GRCh38 as human reference. The output-filtered gene expression matrices were analyzed in *‘R’* (v.4.2.1) with the *‘DropletUtils’* (v.1.8.0) and *‘Seurat’* (v.4.3) packages. In brief, for each sample, cells were detected by ranking cell barcodes according to their number of unique molecular identifiers (UMIs) captured using the barcodeRanks function. Low-ranked cells from this process were labeled as false positives and were discarded, yielding 218^*^10^6^ unique reads with an average of 4000 reads per cell.

The criteria for filtering the dataset were as follows: (i) genes expressed in more than three cells; (ii) cells expressing more than 200 genes; (iii) low-quality cells were removed if they included more than 25% UMIs from the mitochondrial genome. Gene expression matrices were normalized by the NormalizeData function, and 2,000 features with high cell-to-cell variation were calculated using the FindVariableFeatures function. We identified ‘anchors’ between individual datasets with the FindIntegrationAnchors function for batch correction of gene expression of all cells across sequencing datasets. The dimensionality of the dataset was reduced to 100 principal components of the linearly scaled data using the ScaleData and RunPCA functions, respectively. Finally, we clustered cells using the FindNeighbors and FindClusters functions and performed nonlinear dimensionality reduction by uniform manifold approximation and projection for dimension reduction (UMAP) with the RunUMAP function, using 30 dimensions for all approaches. The FindAllMarkers function in Seurat was used to find markers for each unique cluster. Clusters were identified and annotated based on expression of canonical markers for epithelial cell types (Supplementary Table S1).

### Cell interaction Analysis

To identify and visualize the cell cross-talk among cells or between clusters, the R package ‘CellChat’ (v.1) was used according to the developer’s vignette [https://github.com/sqjin/CellChat] [12]. ‘CellChat’ leverages known ligand-receptor pairings to determine the probability of signaling interactions at a single-cell resolution. Each pathway of the network is determined by summing all interaction strengths of the ligand-receptor pairs, via a quantification of their respective expression levels on different cell types [12].

### Differential Pathway Enrichment Analysis – PROGENy

To identify molecular pathways that are differentially regulated between moderate and severe PCS patients, the PROGENy database was used [13]. PROGENy leverages a combination of publicly available signaling experiments to identify core responsive genes associated with molecular pathways. The total expression of downstream gene targets associated with the respective pathway was collated to determine its enrichment score.

### Determination of cell abundance

The ‘scProportion Test Analysis’ investigated the difference between the proportion of cells in clusters between two scRNA-seq samples. A permutation test was used to calculate a statistical p-value for each cluster, as well as, a confidence interval for the magnitude difference is returned via bootstrapping [14]. Differential abundance analysis was also applied via the input of the algorithm *DAseq* (version 1.0.0). This involved the union of data from moderate and severe PCS after dimension reduction.

### Pseudotime

Single-cell pseudotime trajectories were constructed using Monocle (version 2.6.4) [15]. Briefly, we first selected a set of ordering genes that showed differential expression between the clusters. Subsequently, Monocle then uses reversed graph embedding, a machine learning technique to generate a parsimonious principal graph, and then reduces the given high-dimensional expression profiles to a low-dimensional space. Individual cells are projected onto this space and ordered along trajectories that are connected by branching points, which correspond to cell fate decisions. The resulting graph structure was then characterized for network centrality as a proxy of wild-type or impaired cell differentiation.

### Air-liquid Interface (ALI) culture

Four transwell permeable supports (#PID0738600, Corning) were coated with 1mg/ml collagen IV (#5005, Advanced BioMatrix). Healthy donor nasal epithelial cells (NECs) were collected using an interdental brush with written and informed consent. Collected brushings were stored in BEGM with penicillin/streptomycin in a 15ml falcon tube. After washing with phosphate-buffered saline and a cycle of centrifugation. NECs were seeded in supplemented BEGM media (#CC3171, Lonza) in both the apical and basal chambers. Once confluence was reached, the medium was removed from the apical chamber to initiate cellular differentiation. The media in the basal chamber was changed to PneumaCult ALI maintenance medium (#05006, Stem Cell technologies) with or without 10 ng/ml TNFα (#AB50036, Abcam). All media were supplemented with Pennecilin/Streptomycin (#P0781, Sigma-Aldrich) The medium in the basal chamber was changed every two days. 28 days after initiation of differentiation, cells were covered with a preservation buffer (Singleron, Cologne, Germany) and shipped at 4°C for single-cell sequencing analysis. A schematic of the ALI methodology is in the supplement, see Supplementary Fig. S1.

### Differential Gene Expression and Gene Set Enrichment

Differential gene expression of the single-cell data was calculated per cell cluster and PCS group via a Wilcoxon rank sum test as implemented in the R presto library [16]. Gene set enrichment was calculated using a non-parametric Wilcoxon Mann-Whitney test on the log fold changes between the cell groups as implemented in R library gage (v. 2.52) [17].

### TriNetX cohort selection and analysis

We retrieved a case and control cohort for post-COVID symptoms according to the International Statistical Classification of Diseases and Related Health Problems (ICD-10) codes from the TriNetX Global Collaborative Network [18]. This network provides access to electronic medical records from 107 healthcare organizations (HCOs). This retrospective study is exempt from informed consent. The data reviewed is a secondary analysis of existing data, does not involve intervention or interaction with human subjects, and is de-identified per the de-identification standard defined in Section §164.514(a) of the HIPAA Privacy Rule. The process by which the data is de-identified is attested to through a formal determination by a qualified expert as defined in Section §164.514(b)(1) of the HIPAA Privacy Rule. A total of 77 providers responded with a total of 52,833 samples having a diagnosis of post-COVID syndrome disease, unspecified (U09.9; ICD-10-CM) and must not have COVID-19 (U07.1; ICD-10-CM) and COVID-19 virus not identified (U07.2; ICD-10-CM). As controls, we selected people presenting for general examination with no complaints “Encounter for general examination without complaint, suspected or reported diagnosis” (ICD10CM:Z00) and had neither COVID-19 nor post-COVID diagnosis, resulting in a total of 15,797,934 patients from 105 HCOs. To balance the cohorts, propensity score matching was performed based on age, age at index, and gender to address confounding factors. The selection process is represented in Supplementary Fig. S2. A total of 69 diseases related to respiratory and nasal complications were selected according to their ICS-10 codes (Supplementary Table 1) and their odds and hazard ratios between cases and controls were calculated according to the “Compare Outcomes” function of the TriNetX online interface.

**Table 1:** Summary of patient clinical data. Data is presented as the arithmetic mean. The statistical significance between the groups is calculated by a Chi-Square test and significance (p < 0.05) is indicated by an asterisk (**\***) for mild vs. moderate and a hashtag (**#**) for moderate vs. severe. *SD = standard deviation; ENT = ear, nose, and throat; PCS = post-COVID syndrome*.

|  | Mild | Moderate | Severe |
| --- | --- | --- | --- |
| Sex, n (F/M)* | 0/5 | 6/5 | 11/6 |
| scRNA-seq analysis | 4 | 11 | 14 |
| Age (+/- SD) | 47.40 +/- 10.69 | 37.56 +/- 11.47 | 40.41 +/- 13.81 |
| PCS score (+/- SD) | 3.4 +/- 3.27 | 22.14 +/- 4.69 | 35.71 +/- 5.96 |
| Olfactory dysfunction; % yes # | 60.00 | 54.55 | 64.71 |
| Olfactory and gustatory dysfunction; % yes * | 0.00 | 27.27 | 58.82 |
| Fatigue; % yes*# | 0.00 | 81.82 | 94.12 |
| Lack of physical endurance; % yes*# | 20.00 | 63.64 | 82.35 |
| Joint and muscle pain; % yes*# | 0.00 | 18.18 | 58.82 |
| ENT symptoms; % yes# | 0.00 | 0.00 | 23.53 |
| Breathing problems; % yes # | 0.00 | 0.00 | 29.41 |
| Heart problems; % yes # | 0.00 | 0.00 | 23.53 |
| Gastrointestinal problems; % yes # | 0.00 | 0.00 | 23.53 |
| Memory problems; % yes*# | 40.00 | 90.91 | 100.00 |
| Skin complaints; % yes# | 0.00 | 0.00 | 35.29 |
| Symptoms of infection; % yes*# | 0.00 | 36.36 | 76.47 |
| Sleep difficulties; % yes*# | 0.00 | 90.91 | 94.12 |

## Results

### Post-COVID symptom severity correlates with multi-organ issues and increased risk of respiratory co-morbidities

Nasal biopsies were collected from 33 patients recruited from the NAPKON study presenting with mild (n = 5), moderate (n = 11), and severe (n = 17) post-COVID symptoms (PCS). The nasal epithelium was sampled from the anterior and medial head of the middle turbinate using a curette (Figure 1). All patient groups had a comparable mean age, although we noted a gender imbalance in the mild and severe PCS groups. Patients reported olfactory dysfunction to the same percentage of about 60%, irrespective of their PCS classification. The reported gustatory dysfunction, lack of physical endurance, and muscle pain correlated with PCS severity. Patients with severe PCS additionally suffered from ENT, respiratory, and heart problems (Table 1).

### scRNA-seq analysis identifies differently stratified nasal epithelia with PCS severity

To investigate differences in the cellular composition, pathway activities, and cell-cell communication of the nasal epithelia from the three PCS groups, we performed single-cell transcriptome analysis (scRNA-seq) on the 33 patient samples. After sequencing, four samples were excluded from further analysis due to low cell count or poor quality, as they did not meet the quality control criteria. Supplementary Fig. S3 provides a detailed flow of the sample size throughout the analysis. The remaining 29 samples comprised a total of 56,624 cells, after excluding cells with more than 25% mitochondrial gene content and less than 200 genes. Dimension reduction by Uniform Manifold Approximation And Projection (UMAP) of the gene expression matrix identified 17 unique cell clusters. Cluster annotation was done based on known markers for epithelial cell types in the conducting airways. These consist of keratin 5/7 (KRT5/7) for proliferating and differentiating basal and mucosal cells, tubulin beta 4B class IVb (TUBB4B) for ciliated cells, and cystic fibrosis transmembrane conductance regulator (CFTR) for ionocytes [19]. The annotation confirmed the presence of known nasal epithelial, as well as immune cells (Fig. 2A), such as basal and activated myeloid dendritic and T cells. The former are classified through the expression of CD68, and CD83+, the latter via T cell receptor beta constant 2 (TRBC2). Cells from individual patients were evenly spread in the UMAP plot and showed no sample bias concerning cell type or samples (Supplementary Fig. S4 and Supplementary Table S1 for a complete list of differentially expressed genes and canonical markers for cluster annotation). For subsequent analysis, we reduced the number of cell clusters to 12, collapsing differentiating basal, goblet, ciliated, and proximal ciliated cells to one cluster each, based on the similarity in marker gene expression (Fig. 2B).

**Figure 2:**
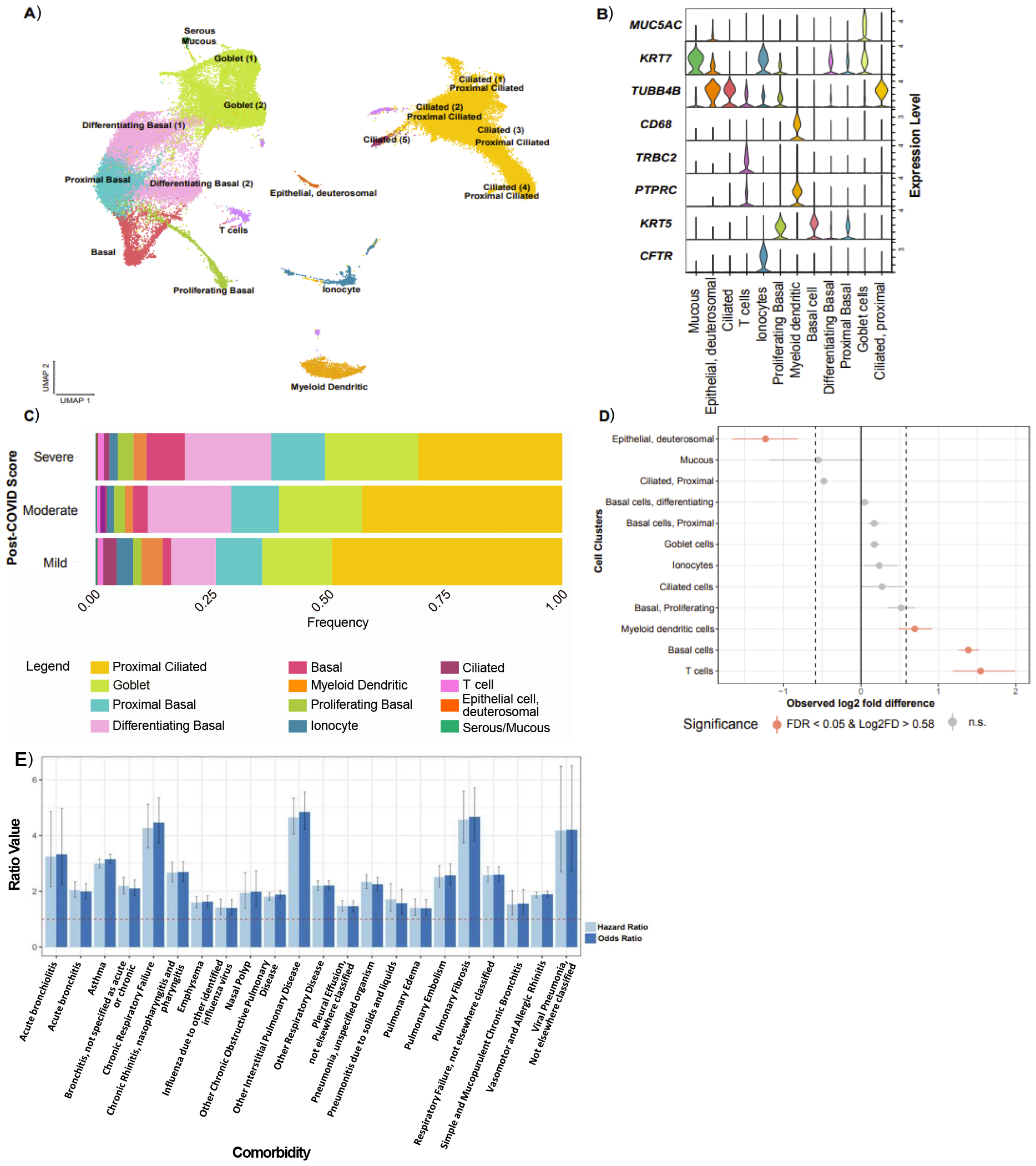
scRNA-seq analysis of the cellular composition of nasal samples from PCS patients. **(A)** Cell type revealing UMAP plot of all integrated samples. 17 distinct cell clusters were detected by cluster gene signatures. (**B**) Violin plots of marker gene expression (*ln*-transformed counts per million) for the 17 distinct clusters collapsed to 12 common cell types found in the conducting airways. **(C**) Stacked bar plots of cell cluster frequency in mild, moderate, and severe PCS patients. **(D)** Relative differences in cell cluster proportions between moderate and severe PCS groups. Red dots indicate statistical significance determined by FDR < 0.05 and a mean log2-fold difference > |0.58| as indicated by the black, dashed lines (permutation test; n=10,000). **(E)** Hazard and odds ratios of 22 nasal and respiratory diseases significantly increased in post-COVID patients. Each ratio is plotted in pairs. Light blue and dark blue indicate the hazard ratio and odds ratio with confidence intervals, respectively.

Previous studies identified the nasal and upper respiratory tract as the primary SARS-CoV-2 infection site, leading to an altered nasal epithelial composition [20]. Accordingly, we first searched for the presence of viral sequences and markers for inflammation in our scRNA-seq data. We neither detected Sars-CoV-2 genetic sequences nor a significant upregulation of *IL6* or *CXCL8 in severe sa*mples, indicating viral clearance and the absence of an active inflammatory response to infection (Supplementary Fig. S5). However, we observed a significant change in the relative airway cell composition that correlated with PCS severity (Fig. 2C): proximal ciliated cells and their precursors, deuterosomal epithelial cells, decreased in relative abundance with increasing PCS score. Quantifying the changes via permutation testing and bootstrapping [21], indicated a 0.5 log2-fold reduction of both proximal ciliated and mucous cells and a statistically significant reduction of deuterosomal epithelial cells (1.2 log2 fold change, p<0.05) in severe compared to moderate PCS (Fig. 2D). Conversely, the abundance of T-, basal and myeloid dendritic cells significantly increased, respectively, 1.5, 1.3 and 0.6 log2-fold with severe PCS. Changes in cell type abundance were independently confirmed by the *DA-seq* method, which detect differential cell population abundance using a *neighborhood-based* multiscale measure. In severe PCS, the algorithm found ciliated cells to be depleted and basal epithelium, T cells, and myeloid dendritic cells as enriched compared to moderate PCS (Supplementary Fig. S6).

With a deficiency of ciliated and goblet cells and epithelial mis-differentiation in PCS, we hypothesize that the projective function of the nasal epithelium against pathogens and irritants is diminished. To check whether this condition increases the risk for infections through the respiratory tract, we extracted diagnosis information from 107 healthcare organizations worldwide using the TriNetX research network [18]. A search for patients with post-COVID-19 disease according to ICD-10 codes (Supplementary Table S2) returned 52,833 cases and 51,310 propensity-matched controls out of almost 130 million patients (Fig. 2E). Screening for 69 respiratory and nasal comorbidities, 22 diseases showed significantly increased odds and hazard ratios (OR and HR > 1; Supplementary Table S2). Among the highest comorbidities in these patients were chronic respiratory failure, interstitial pulmonary disease, pulmonary fibrosis, and viral pneumonia (OR and HR > 4), but also acute bronchiolitis and asthma (OR and HR > 3).

Our scRNA-seq analysis thus confirms a lasting, possibly even worsened, tendency towards altered stratification of the nasal mucosa that correlates with PCS severity, even in the absence of any viral infection. The number of undifferentiated basal cells is increased at the expense of reduced numbers of ciliated cells. Consequently, there is a higher presence of immune cells resident in the nasal epithelium. PCS is accompanied by an increased risk for respiratory disease, which might be linked to the reduced barrier function against external pathogens due to mis-differentiation.

### Aberrant nasal epithelial composition is driven by TNFα and TGFβ signaling in the immune cell compartment, affecting basal epithelial cell differentiation

To investigate the signaling pathways responsible for the differences in epithelial stratification between moderate and severe PCS, we first examined the enrichment of upstream pathways leveraging the PROGENy [13] pathway compendium. Severe PCS samples indicated enrichment in inflammation-related pathways (TNFα, TGFβ, NF-κB in Fig. 3A). This, combined with an increased abundance of myeloid and T cells (Fig. 2D) is indicative of a more activated immune response in severe versus moderate PCS. Interestingly, the PI3K pathway is more enriched in moderate PCS samples, which could have different implications. This observation might indicate a reduction in cell proliferation processes or a preference for SMAD signaling and proliferation over PI3K/mTOR-controlled cellular migration in cases of severe PCS.

**Figure 3:**
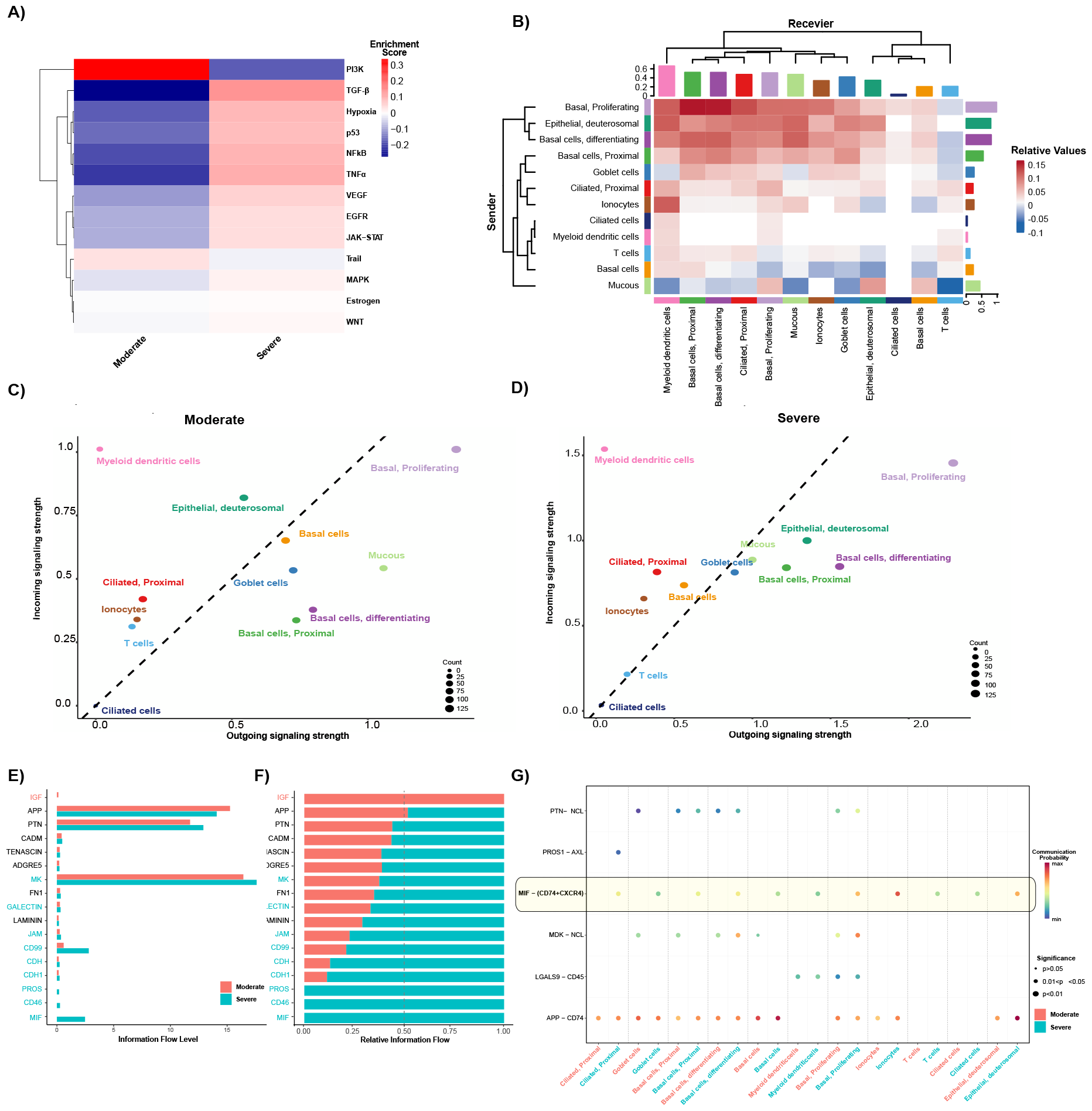
Enriched pathological pathways, and cell-cell interactions with PCS severity. (A) Relative pathway enrichment in moderate and severe PCS. Red indicates positive, whilst blue indicates negative enrichment. (B) Heatmap of the signaling pathway enrichment contributing to outgoing or incoming communication. The color bar indicates the relative signaling strength in PCS severity; red = increased in severe PCS and blue = decreased in severe PCS. The solid-colored bars on the x and y axes indicate the sum of the incoming (x-axis) or outgoing (y-axis) signaling strength for each cell type. Comparison of total incoming signaling strength vs. total outgoing signaling strength across cell populations in moderate (C) and severe (D) PCS. All significant pathways (accumulated p-value <0.05) presented as absolute (E) or relative (F) information flow, ranked based on differences in the overall information flow between moderate (red) and severe (cyan) patients. (G) Representation of significant ligand-receptor pair interactions from all other cell groups to myeloid dendritic cells in moderate (red) and severe (cyan) PCS. The dot color and size represent the calculated communication probability and p-values, respectively.

The restitution of a functional nasal epithelium is regulated by carefully orchestrated incoming and outgoing cell-cell communication networks. Using *CellChat* [12], we inferred the strength, direction, and relative change with PCS severity via the differential expression of known receptor-ligand gene pairs. We observed in moderate PCS basal proliferating cells are identified as sending the most molecular signals, whilst myeloid dendritic cells are enriched as receivers of incoming signals (Fig. 3B - 3D), in both moderate and severe PCS. These three cell types appear to form a cell communication nexus in severe PCS (Fig. 3B, top left). Comparatively, in moderate PCS, mucous and basal cells send, and T cells receive more signals (Fig. 3B, bottom right). Myeloid dendritic cells receive input from all cell types, irrespective of PCS severity, but specifically send signals to basal proliferating and T cells in severe PCS (Figs. 3C-G).

Clearly, important changes in cell-cell communication networks exist with worsening PCS. We next analyzed the differential signaling strengths of individual pathways between moderate and severe PCS. Of note, extracellular matrix (ECM) and cell receptor pathways, although involved in PCS, remained unchanged between moderate and severe groups (tenascin, laminin, or fibronectin), amyloid precursor protein (APP) and cell adhesion molecule (CADM) interactions (Figs. 3E and 3F). Interestingly, the IGF pathway (related to cell differentiation [22]), is absent in severe PCS, corroborating with our previous observation of a reduction in differentiated epithelial cells (Fig. 3E). Instead, cell-cell communication in severe PCS is dominated by cell-cell adhesion (CD46, CD99, JAM), immune signaling (GALECTIN) and growth and survival (MK, PROS1). The macrophage migration inhibitory factor (MIF) demonstrated the strongest enrichment in ligand-receptor interactions in severe PCS. MIF is an inflammatory cytokine, linked to TNFα and TGFβ [23, 24]. MIF signalling via the CD74, CXCR4, and CD44 receptors activate downstream NF-κB, MAPK, and AKT pathways, which regulate immune responses, inflammation, proliferation, and cell survival depending on the receptor combination [25, 26]. *MIF* and *CD74* are both highly expressed by all cell types, irrespective of PCS severity (Supplementary Figs. S7A, B). We observe that *CXCR4* (myeloid) and *CD44* (myeloid and basal cells) indicate increasing expression with PCS severity (Supplementary Figs. S7C, D). Accordingly, *CellChat* analysis reveals specific enrichment for the MIF-(CD74-CXCR4) stimulatory axis in myeloid cells in severe PCS (Fig. 3G), whilst basal proliferating cells indicate an enrichment of signals received via CD44 in severe PCS patients (Supplementary Fig. S8). The enrichment of these receptor-ligand interactions in a severity-specific manner indicates the activation of potentially distinct pathways between the PCS groups contributing to the observed phenotype.

To elucidate the correlation between paracrine cell-cell communication with PCS severity, we inferred the upstream pathways per cell type using PROGENy, i.e., linking gene expression to ligand activation. The analysis revealed an enrichment of EGFR, and NF-κB pathways in both myeloid dendritic and T cells (Fig. 4A), confirming the downstream effect of CD74 activation. Also, PROGENy predicted upstream activity of TNFα, possibly due to the significant overlap in NF-κB and TNFα target genes. Because of the upstream signaling, we find increased expression of *TNF* and *TGFB1* in myeloid and T cells with PCS severity (Fig. 4B). In alignment, *TNF* expression is observed primarily in the immune cell compartments, with this expression likely driven by upstream activation of the NF-κB pathway. *TGFB1* expression increases in both immune and basal epithelial cells (Fig. 4B). Its upregulation possibly stems from two different upstream causes. In the former cells, it is putatively caused by upstream TGFβ signaling, while in the latter through WNT or MAPK pathway activation (Figs. 4A and 4B). Interestingly, in myeloid cells there has been an inhibitory action of TGFβ towards TNF observed, causing cancer growth, and also priming myeloid cells to osteoclastogenesis, an inflammation phenotype associated with rheumatoid arthritis [27-29]. As a result of TNFα and TGFβ1 secretion, all cell types, but mostly basal epithelial cells, responded with an increase in *TNF and TGFβ* receptor expression (Supplementary Fig. S9). Both TGFβ receptors, TGFβR2 and TGFβR1 work together as a receptor complex to transmit signals from TGF-β ligands to the intracellular machinery [30].

**Figure 4:**
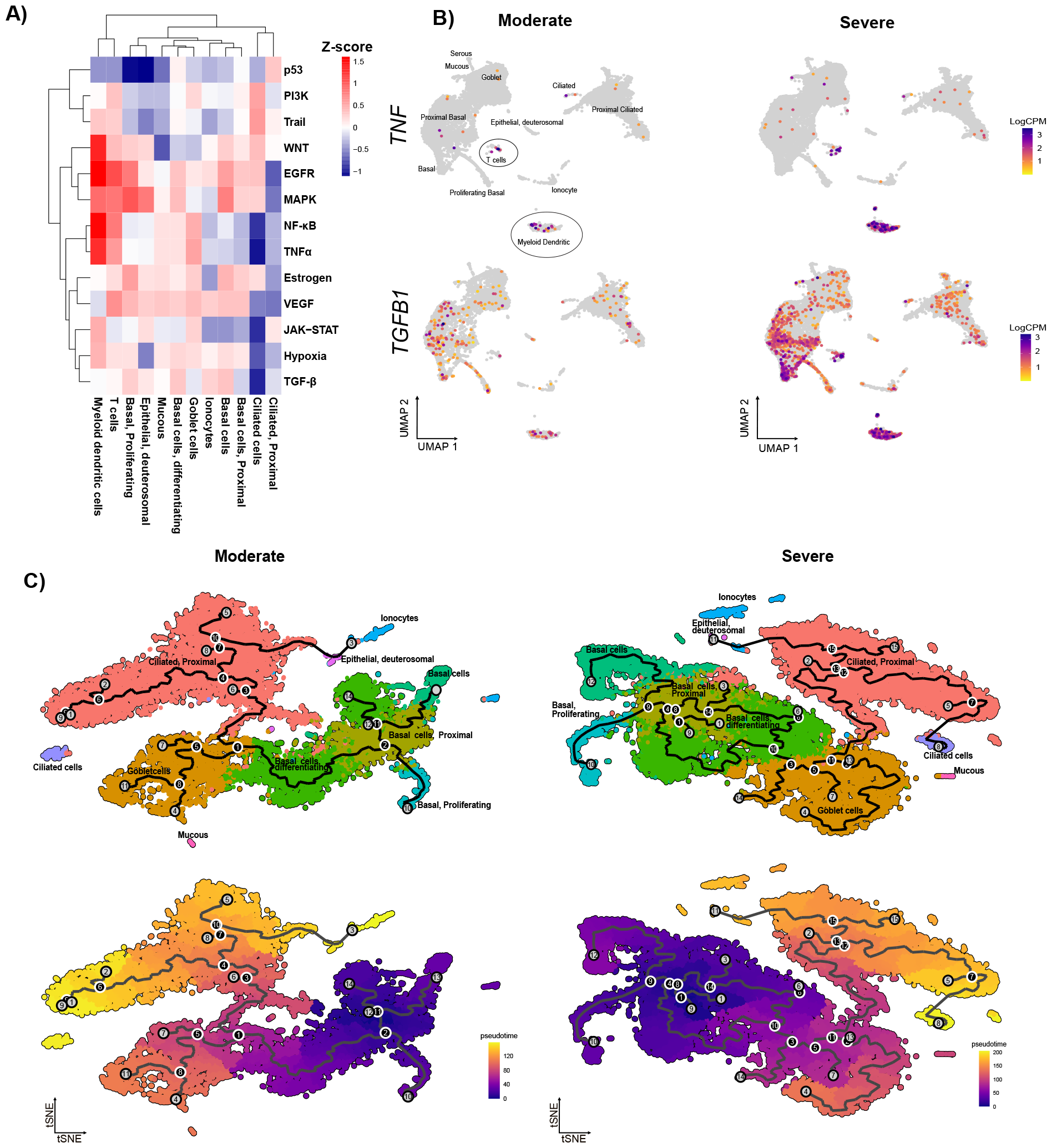
Presentation of cellular expression of *TNFα* and *TGFβ* and altered basal cell differentiation trajectory. (A) Heatmap of Progeny pathway enrichment, stratified by cell type; blue = enriched in moderate PCS and red = enriched in severe PCS. (B) UMAP of *TGFB1, TGFB2*, and *TNF* expression; purple = increased expression, and grey = no expression. Gene expression is quantified as log2CPM. (C – F) Pseudo-time analysis of nasal epithelial cell differentiation in moderate (C & D) and severe (E & F) PCS patients. (C & E) t-SNE of cell populations. (D & F) t-SNE plot with overlaid of pseudo-time progression of cell development; purple = earlier in trajectory and yellow = later in trajectory. Black lines represent the most likely path of cell maturation over the pseudo-time trajectory.

TGFβ signaling is linked to Epithelial-Mesenchymal Transition (EMT), in which alterations of epithelial cells lead to mesenchymal characteristics. EMT is an important factor in remodeling of the nasal epithelium during inflammation [31]. To check whether TGFβ signaling leads to EMT, we performed a Gene Set Enrichment Analysis (GSEA) [32] on the gene expression log fold changes of basal epithelial cells between mild and severe cases using 2760 and 12362 cells, respectively. Using the human Hallmark Gene set [32] we find indeed TNFα, NF-κB as well as EMT as the most upregulated pathways in the basal epithelial cells of severe PCS patients (Supplementary Fig. S10).

In summary, our analysis shows that basal cells and dendritic myeloid cells are integral communicator cells in PCS. However, in severe PCS, our data shows that the profile of these cells shifts with basal proliferating cells producing a TGFβ-driven response and myeloid cells maintaining an inflammatory profile. Inflammation is triggered through MIF and subsequent TNFα signaling, leading to the activation of TGFβ and EMT in the post-COVID nasal epithelium.

To confirm our hypothesis of incorrect EMT and epithelial differentiation, we used pseudo-time trajectory analysis across both PCS severity. Pseudo-time analysis positions cells along a trajectory that quantifies the relative progression of differentiation. Trajectories through the high-dimensional expression space can be complex, with branches to multiple endpoints. Accordingly, Figures 4C–4F depict the trajectories of nasal epithelial cell differentiation within moderate and severe PCS patients on t-SNE plots. The number of edges for both differentiation trajectories in severe and moderate PCS are 327 and 247, respectively. Notably, these data indicate a more branched pseudo-time trajectory with multiple endpoints for severe PCS, which is reflected in the lower degree of centrality of 8.1×10^−3^ vs 3.1×10^−3^ in moderate and severe PCS. Thus, there is indeed a divergence from the normal differentiation route from basal epithelial differentiation to ciliated cells with severe PCS.

### TNFα and TGFβ exposure causes aberrant differentiation of basal epithelial cells

Single-cell transcriptomics of our PCS samples revealed an upregulation of TNFα and TGFβ signaling through the presence of immune cells, which potentially promotes EMT and aberrantly stratified nasal epithelium. We report that the primary epithelial cells stimulated by TNFα are differentiating basal and goblet cells in severe PCS (Fig. 4A). As such, to validate whether TNFα is a causal factor promoting a bias against the formation of ciliated cells, we differentiated nasal epithelial cells *in vitro* using an air-liquid interface (ALI) model with or without TNFα stimulation (Supplementary Figure S1). 28 days after initiation of differentiation, cells underwent single-cell sequencing analysis. A UMAP visualization, clustering, and annotation using the Human Cell Atlas revealed the presence of seven distinct cell clusters (Fig. 5A), wherein we identified ciliated cells based on *Tubb4B* expression (Cluster 5, Figure 5B) and basal cells in clusters 2-4 based on KRT5 expression.

**Figure 5:**
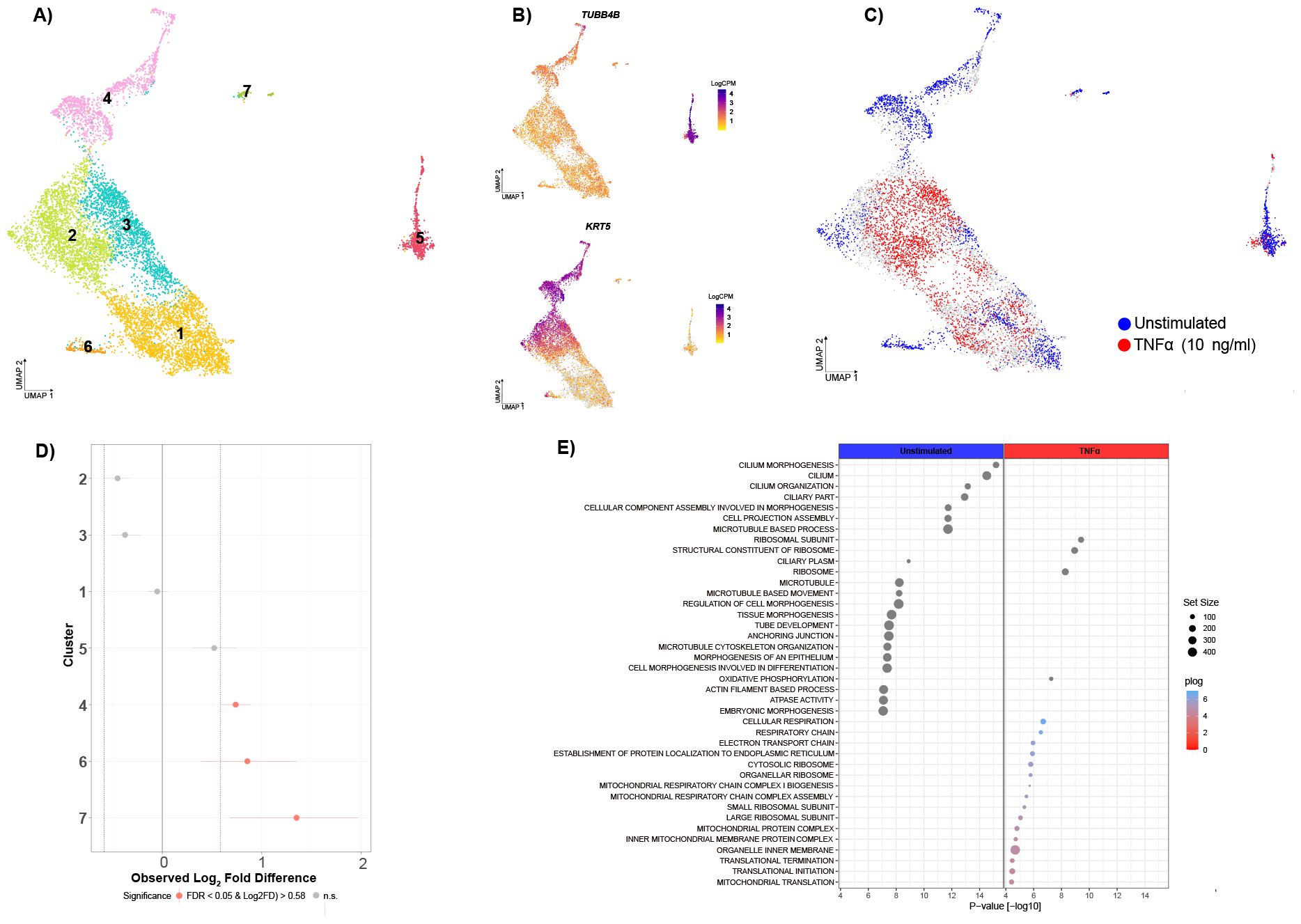
scRNA-seq analysis of ALI cultures stimulated with TNFα. (A) UMAP of integrated scRNA-seq of primary nasal epithelial cells differentiated over 28 days with and without TNFα (10 ng/mL) stimulation. Expression of *TUBB4B* (B) and *KRT5* (C) to identify ciliated and basal cell locality in UMAP, respectively. (D) Differential abundance analysis of cell groups by stimulation conditions, red = increased in TNFα-stimulation and blue = increased in unstimulated cells. (E) Relative differences in cell proportions for each cluster between the TNFα-stimulated and unstimulated conditions. Clusters colored red have an FDR < 0.05 and mean Log2 fold difference > |0.58| compared to the severe patients (permutation test; n=10,000). (F) Pathway enrichment analysis stratified by treatment conditions. Left/red presents cells enriched in TNFα-stimulated cells and right/blue presents pathways enriched in unstimulated cells; n = 4 for all analysis.

A differential abundance analysis (DA-seq) between the TNFα stimulated, and unstimulated controls revealed a lower abundance of cell clusters 4-7 under TNFα stimulation [0.5, 0.7, 0.8, 1.3 log_2_ fold down-regulation, scProportionTest (Figure 5D). The results propose a decrease in ciliated cells (cluster 5) and loss of epithelial cell diversity in clusters 4-7. To identify signaling pathways included by TNFα stimulation, we performed a GSEA on the bulk-aggregated data using the REACTOME. Stimulation down-regulated pathways related to ciliation, cilium, and differentiation, while it upregulated ribosome and mitochondria-related signaling. Taken together, in our model, TNFα significantly impaired the *in vitro* differentiation of NECs, clearly supporting the results from the PCS patient samples. This implicates increasing TNFα production from the immune cells with PCS severity, affecting normal basal epithelial differentiation and restitution of a normal epithelial barrier.

## Discussion

The Post-COVID syndrome (PCS), is a multisystemic condition often characterized by severe symptoms that persist following an infection with SARS-CoV-2. The global prevalence of the post-COVID syndrome is staggering, with at least 65 million individuals affected worldwide. This number continues to rise daily, based on a conservative estimated incidence rate of 10% among infected individuals and more than 651 million documented COVID-19 cases worldwide. Although, the actual number is likely much higher due to many undocumented cases.

The incidence of post-COVID syndrome varies across different patient populations, estimated at 10– 30% among non-hospitalized cases, 50–70% among hospitalized cases, and 10–12% among vaccinated cases, however clear diagnostic criteria are still lacking making precise estimates difficult [33, 34]. PCS affects individuals of all ages, with the highest percentage of diagnoses typically observed between the ages of 36 and 50 years, with a noteworthy observation that the relative frequency of PCS cases is highest among in non-hospitalized patients [9, 35]. Thus, although the COVID-19 pandemic is diminishing, health concerns are far from over [36]. While a plethora of PCS symptoms have been identified, impacting multiple organs and tissues [9], the mechanisms leading to PCS pathogenesis are still unclear. Proposed mechanisms include immune dysregulation [37, 38], microbiota disruption [39, 40], autoimmunity [41, 42], or clotting through endothelial dysfunction [43, 44]. Here, we have performed a single-cell transcriptome analysis of nasal epithelial biopsies from a cohort of 33 patients with mild, moderate, and severe PCS. As acute SARS-CoV-2 infections are known to cause the destruction of ciliated cells [20], we hypothesized a long-lasting detrimental effect in this tissue even post-clearance of the viral infection.

From our data, we observed a lack of reformation of ciliated epithelial cells in post-COVID syndrome-diagnosed patients, with the degree of mis-differentiation in the nasal epithelium correlating with disease severity. In addition, a prolonged presence of myeloid and T cells is associated with a persistent inflammatory response via MIF-CD74 signaling and increased secretion of TNFα and TGFβ, perpetuating an inflammatory response and epithelial-mesenchymal transition in the basal epithelial cells (summarized in Supplementary Figure S11). We confirmed our *in vivo* predictions that early and sustained exposure to TNFα causes malformation of a pseudostratified epithelium characteristic of the conducting airways through an air-liquid interface model of nasal epithelial cells. Constant exposure to TNFα significantly reduced the number of terminally differentiated ciliated cells and reduced the diversity of epithelial cell types. This novel mechanism for PCS might explain the multitude of nasal and respiratory comorbidities accompanying the conditions, as highlighted in the TrinetX study. A total of 22 diseases show an increased odd/hazard ratio > 1, such as asthma, pulmonary edema, and chronic respiratory failure. Thus, the persistence of infection-like symptoms combined with a poorly restituted respiratory epithelium (identified in the nose) enables recurring infections and disease. These factors possibly promote a positive feedback loop of constant immune cell activation and aberrant nasal epithelium culminating in a reduced protective effect.

Our findings are in line with previous findings of dendritic cells (DCs) and monocytes to be increased in blood for more than 6 months post-severe COVID-19 infections [45]. Roukens et al. detected CD8+ T cells persisting in nasal mucosa for at least 2 months after viral clearance from a SARS-CoV-2 infection [46]. A recent study by Aschmann et al. has linked post-COVID exercise intolerance to artery remodeling in skeletal muscle accompanied by resident immune signatures despite viral clearance. They found a similar long-lasting tissue remodeling and immune response effect in yet another tissue [47]. Apparently, immune cell clearance in the nasal epithelium is impeded even longer in PCS and potentially prolonged by positive feedback through inflammation-promoting paracrine cell-cell communication axis via MIF-CD74 and subsequent production of TNFα and TGFβ. Activation of MIF-CD74 was found to confer protection against oxidative stress in animal models. Genetic silencing of MIF and CD74, as well as pharmacological inhibition of CD74, resulted in the loss of this protective effect, leading to increased inflammatory cytokine production, apoptosis, and higher mortality rates [48]. On a molecular level, CD74 activation during hyperoxia-induced proliferative and pro-survival responses through ERK and Akt pathway activation [49]. The key function of CD74 in stabilizing the presentation of antigens for T cells [23], combined with MIF likely drives the expansion and persistent activation of pro-inflammatory pathways in severe PCS [50]. The persistent production of TGFβ potentially promotes the continued activation of the MIF-CD74 axis, via the ERK pathway, generating a self-perpetuating cycle in severe PCS patients [24].

Cell-cell interaction analysis revealed a complex communication network between primarily basal epithelial, proliferating basal and myeloid cells. In particular, we observed that two MIF co-receptors, CXCR4 and CD44, are enriched for activation in severe PCS. Both receptors are associated with the induction of differential MIF responses in a cell-type-specific manner, but are closely linked to the mediation of immune cell infiltration [49, 51]. In fact, activation of CD74, CXCR4, and CD44 has been associated with COVID-19 and acute lung injury through mediation of the inflammatory immune response [51, 52]. This implicates the appearance or formation of a new immune cell subset in severe PCS potentially contributing to a worsening disease state.

In addition, there is a distinct shift in the interaction between the proliferating basal cell with the ECM proteins laminin B1 and fibronectin. Fibronectin, which is induced by TGFβ stimulation, is well described to induce epithelial proliferation and differentiation [53] and was found enriched in severe PCS. Conversely, laminin B1, which is produced more by basal epithelial cells [54] is enriched in moderate PCS. The laminin protein family has been implicated in modulating correct lung development as well as cell adhesion, migration, and proliferation of epithelial cells [55, 56]. These ECM factors combined with a hypothesized increased deposition of ECM-bound TGFβ produced by basal epithelial cells in severe PCS [57], actions on the proliferating basal cells. Whilst a potentially localized exposure of differentiating basal cells to TNFα from myeloid and T cells promotes reduced formation of ciliated cells, as indicated by our cellular model. The apparent reduced activation of the PI3K pathway in severe PCS, which is associated with reduced cell migration, may result in anoikis (luminal cell extrusion) [58]. This is driven through both ECM remodeling and a ‘crowding’ of basal epithelial cells resulting in a limited delivery of terminally differentiated cells such as ciliated or goblet. As such, PCS comprises an amalgam of structural and cellular interactions at different stages of epithelial formation, which all contribute to the aberrant restitution of correct airway epithelium. This may be indicative of systemic restorative problems in these individuals in their recovery from an acute SARS-CoV-2 infection, contributing to PCS severity.

What remains is the pressing question regarding the resolution of and cure for PCS. Interestingly, chronic inflammation in nasal tissue has been observed in other diseases, such as chronic or allergic rhinitis. Treatment of mice with the TNFα inhibitor infliximab reduced cytokine production and immune cell infiltration into the nasal mucosa [59]. In one case study, pirfenidone treatment was successful in treating post-COVID pulmonary fibrosis [60]. Pirfenidone has anti-inflammatory and fibrotic effects and is used to treat idiopathic pulmonary fibrosis (IPF). While its mechanism of action is not completely understood, it is believed to inhibit collagen synthesis and most importantly suppress the production of pro-inflammatory molecules such as tumor necrosis factor-alpha (TNFα) and transforming growth factor-beta (TGFβ). Moreover, it was found to negatively regulate SMAD and Jak-STAT3 pathways, which are downstream of TGFβ and parallel to PI3K/mTOR signaling [61]. Due to these actions, it was recently suggested as a treatment option for rheumatoid arthritis [62]. Another drug candidate for treatment could be metformin. It has been identified as a repurposed drug for atrial fibrillation (AF) through in silico prediction and *in vitro* testing. AF is characterized by high TGFβ and TNFα levels in serum, with metformin reducing TGFβ production significantly in cardiomyocytes [63]. A recent review highlights metformin’s potential in reducing inflammation (including TNFα and TGFβ), whereby it improved outcomes in infectious diseases, such as influenza or hepatitis-C [64]. Metformin therapy in diabetes was reported to reduce mortality by roughly 70% after SARS-CoV-2 infection [65] and was suggested as a treatment for SARS-CoV-2 infections previously [66].

In conclusion, our findings highlight a potential mechanism for the persistence of PCS long after viral infection. Using single-cell analysis, we detected a deficiency of ciliated cells, heightened immune cell presence, and enrichment of inflammatory pathways in cases of severe post-COVID syndrome in the nasal tissue. It is noteworthy that this cellular and molecular response occurs in the absence of detectable SARS-CoV-2 viral RNA. This contradicts one prevailing hypothesis that PCS is driven by the persistence of a residual viral load [67, 68]. However, this may be due to the sampling location with the potential for some viral load persistence in the lower respiratory tract [69]. Nonetheless, our results indicate that the persistence of infection-like symptoms is driven primarily by an aberrant immune-cytokine-epithelial response axis and opens novel treatment regimens akin to other chronic inflammatory respiratory diseases.

## Notes

### Competing Interest Statement

The authors have declared no competing interest.

### Summary of Updates

The authors included a graphical abstract as a supplementary figure.

