## Supplementary Figures for "scRNA-seq reveals persistent aberrant differentiation of nasal epithelium driven by TNFα and TGFβ in post-COVID syndrome": PCS supplementary figures_17012024.docx

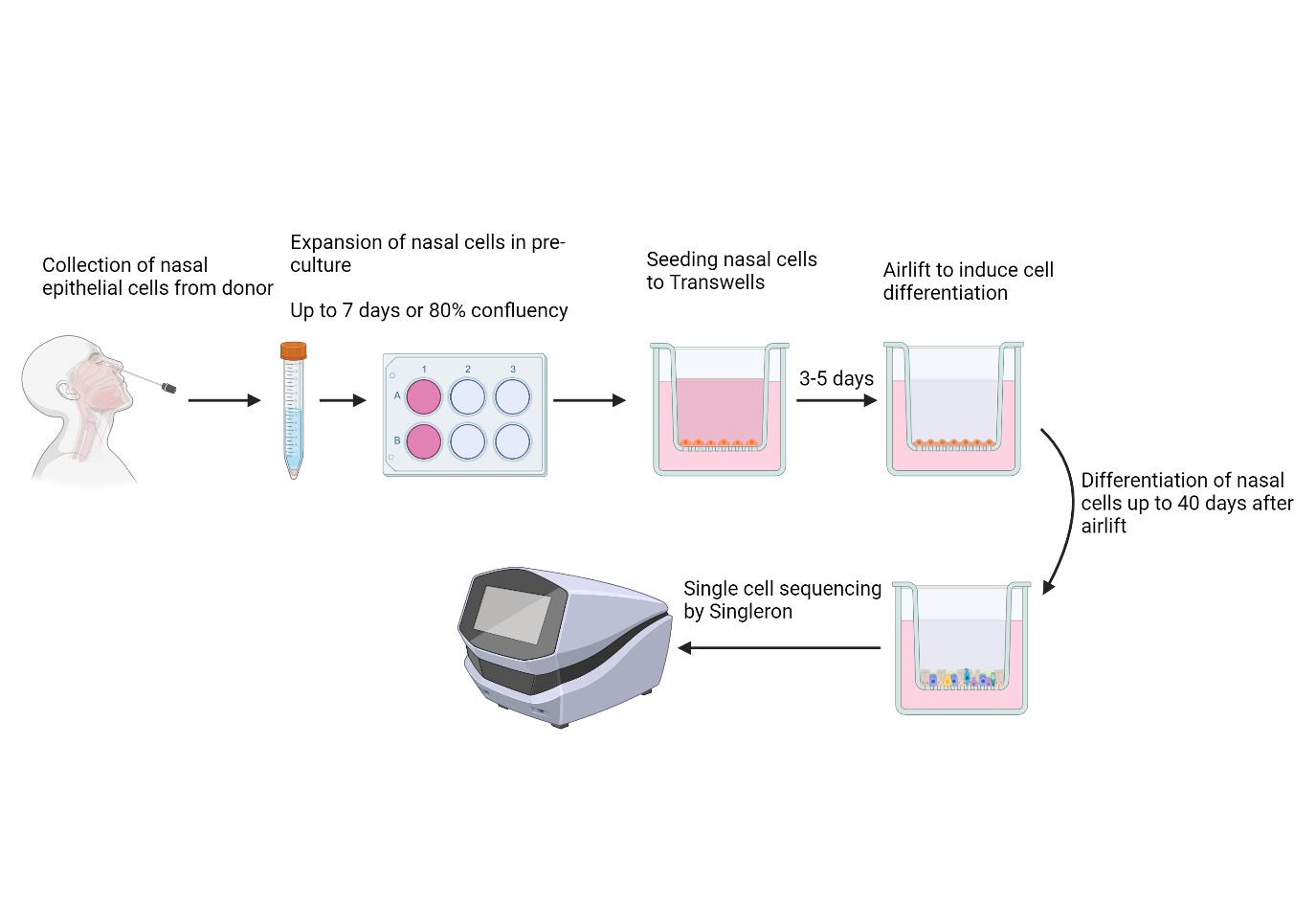

**Figure S1: Schematic of the sample collection, processing, and analysis of air-liquid-interface differentiation of nasal epithelium.**

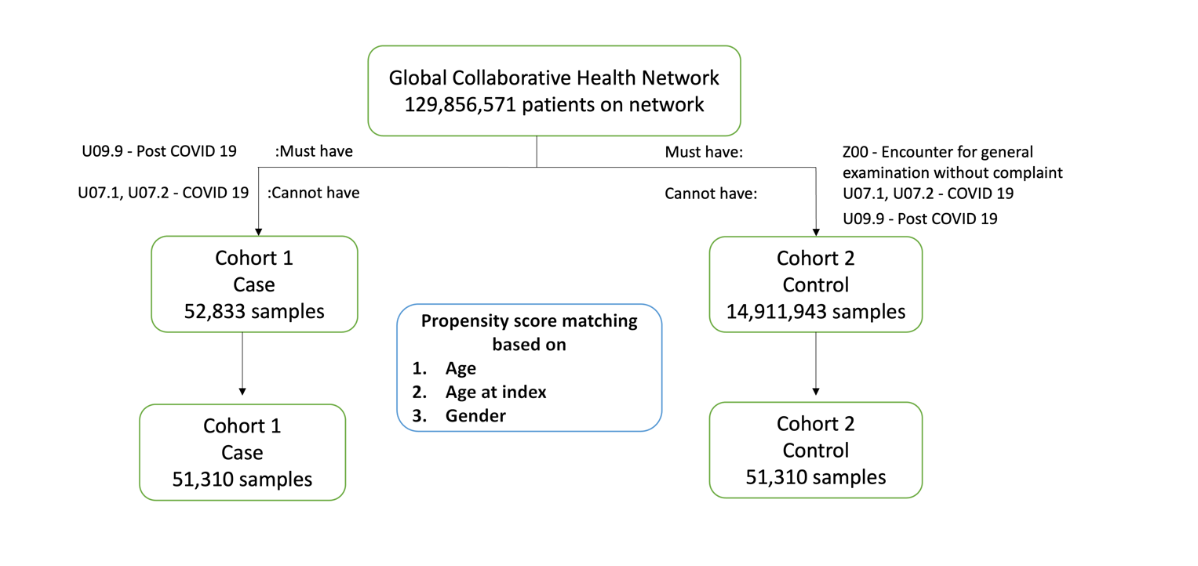

**Figure S2: Outline of the TriNetX database patient selection criteria and generation of analysis cohorts.**

NAPKON

-

POP

n

= 1270

Ca

ses

n

=

33

single

cell

ng

sequenci

n

=

33

n

4

=

Removed for poor sample quality

n = 4

final sample

size

n

=

29

**Figure S3: Outline of NAPKON population samples selected for clinical analysis and single-cell RNA-sequencing.**

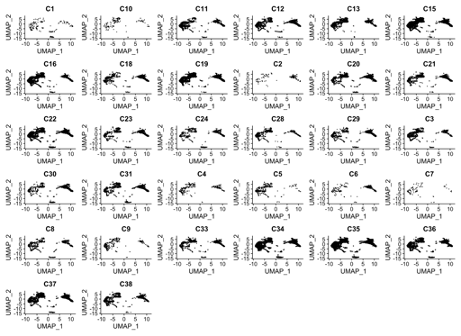

**Figure S4: UMAP illustration of cell populations across all 32 samples sent for scRNA-seq.**

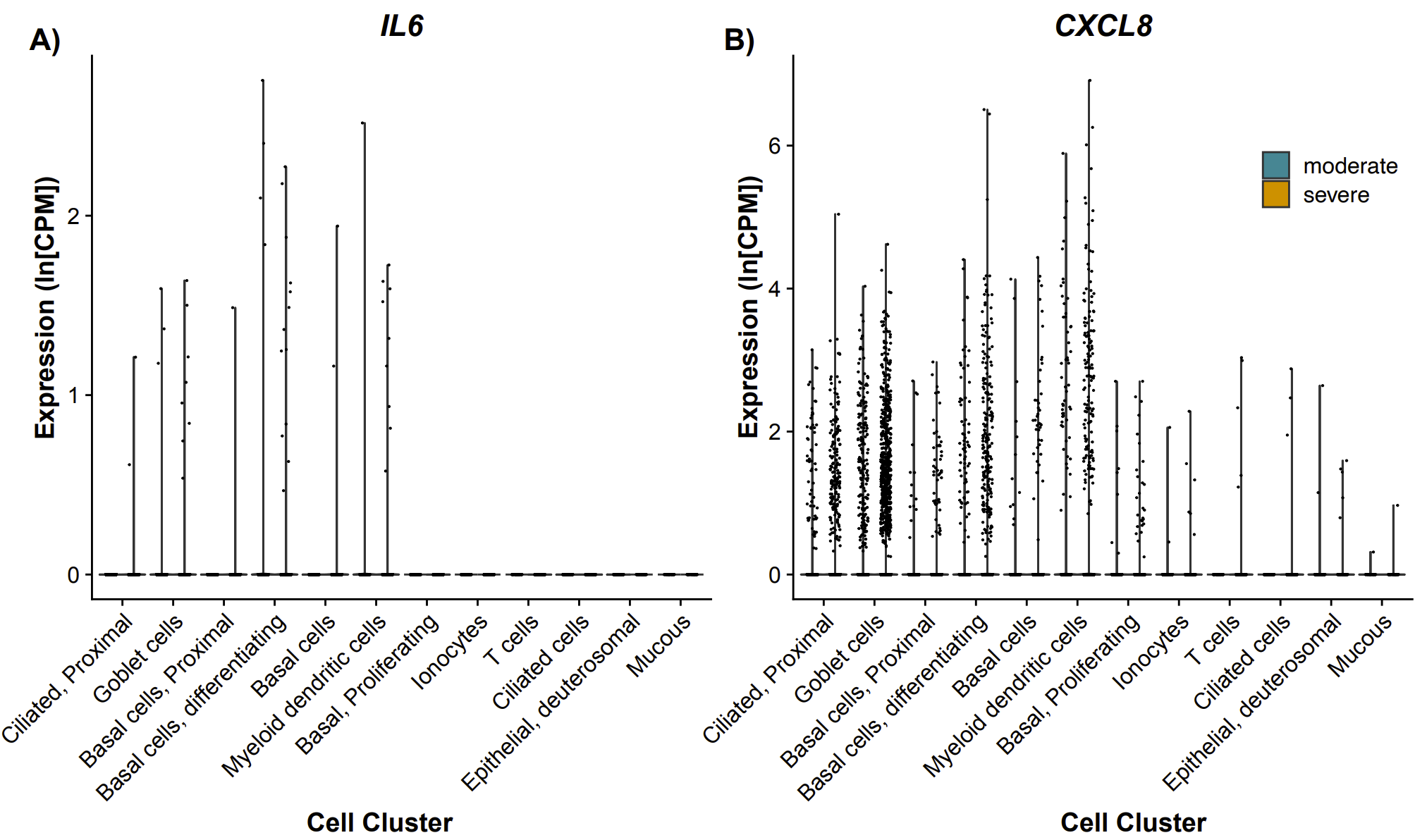

**Figure S5: Gene expression presented as ln(CPM) for *IL6* (A) and *CXCL8* (B) across all annotated cell types.** Moderate PCS patients are indicated in blue (left) and severe PCS in yellow (right). Each point represents an individual cell; n = 29.

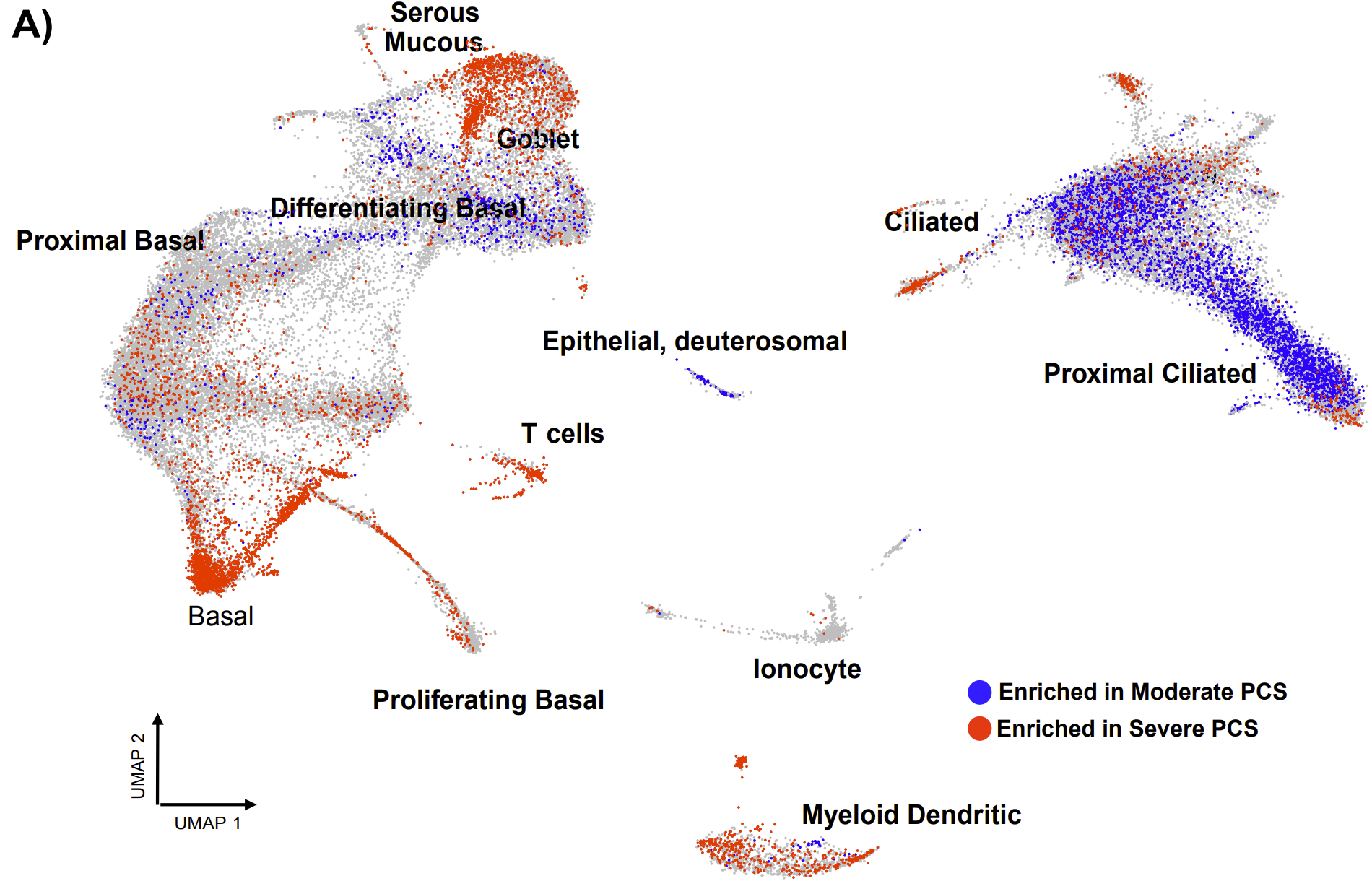

**Figure S6: UMAP representation of differential cell abundance between moderate (blue) and severe (red) PCS.** DA-seq detected differentially abundant cell subpopulations by analyzing cells from moderate and severe PCS. Cells colored blue are more abundant in moderate PCS, whilst red cells are more present in severe PCS, with grey cells equally abundant in both groups. Each point represents an individual cell; n = 29.

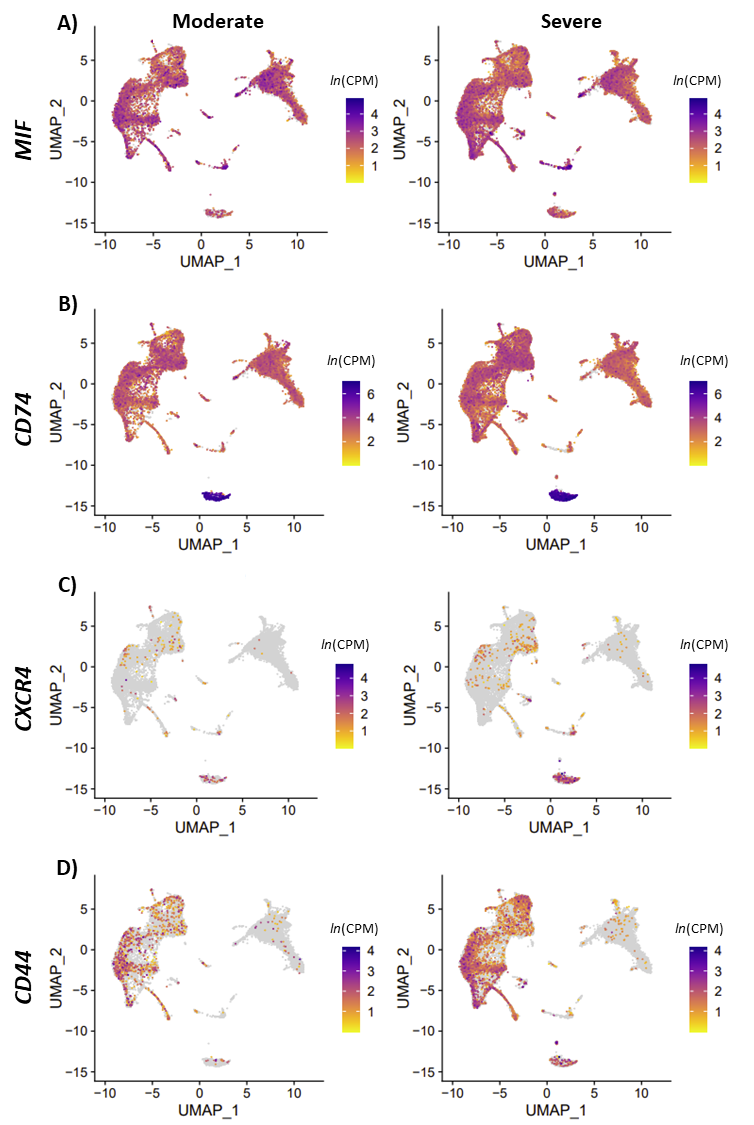

**Figure S7: Gene expression presented on a UMAP for *MIF* (A), *CD74* (B), *CXCR4* (C) and *CD44* (D).** Gene expression is presented as ln(counts per million). Yellow indicates lower expression whilst purple indicates higher expression. The left column presents moderate PCS patients and the right column presents severe PCS patients. Each point represents individual cells; n = 29.

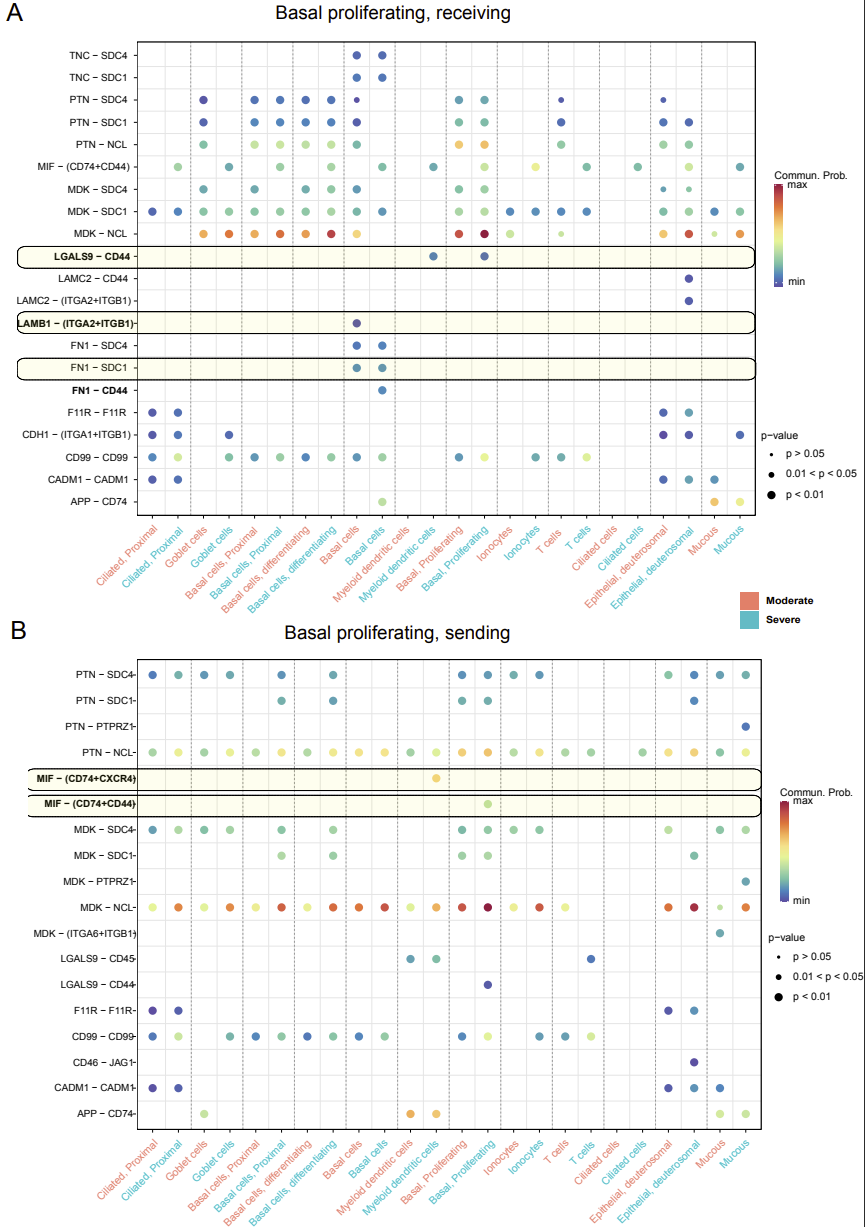

**)**

**)**

**Figure S8: Representation of significant ligand-receptor pair interactions with basal proliferating cells.** (A) Basal proliferating cells are receivers of signals from all other cell types. (B) Basal proliferating cells are senders of signals to all other cell types. The x-axis presents all cell types in moderate (pink) and severe (blue) PCS patients. Statistical significance was determined at p < 0.05; n = 29.

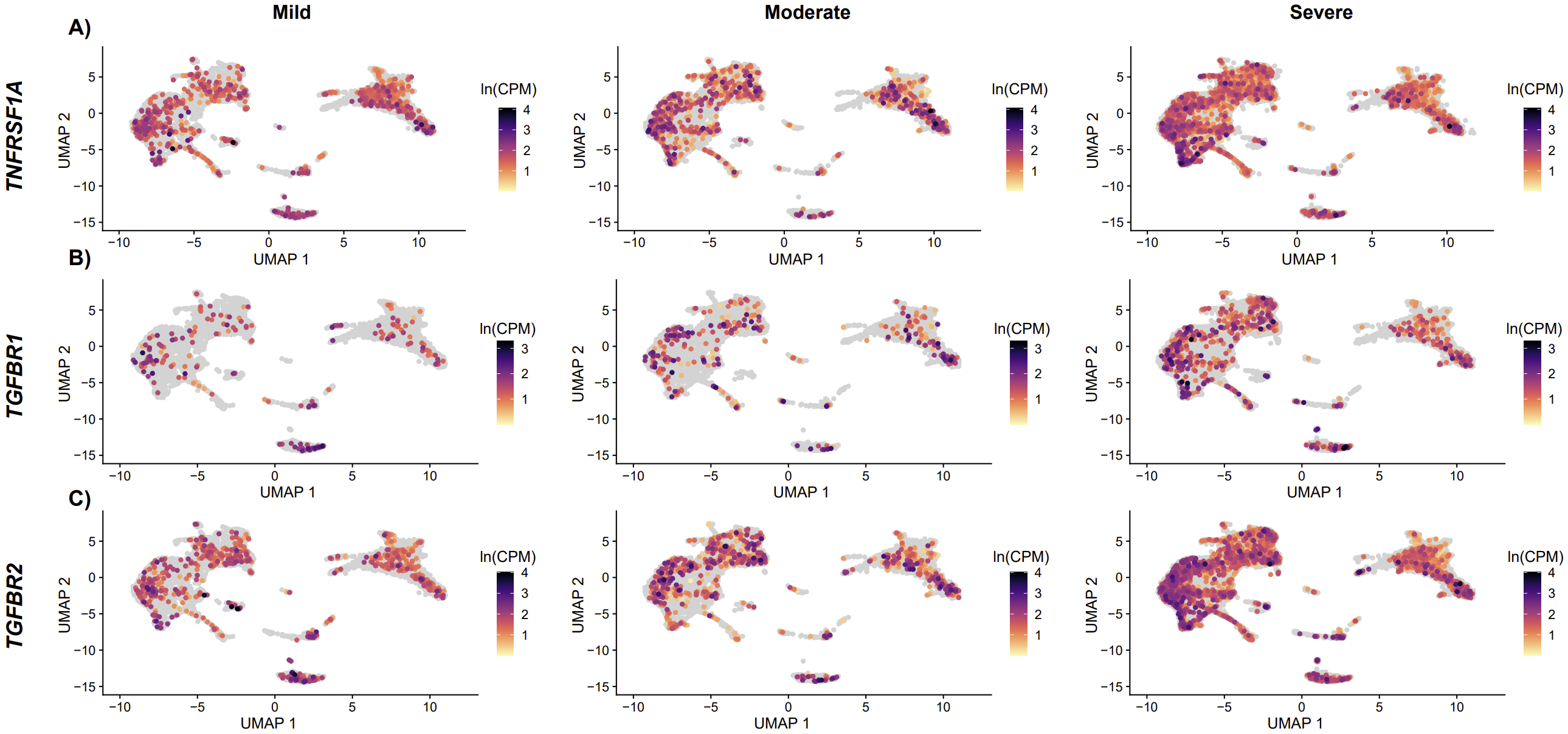

**Figure S9: Gene expression presented on a UMAP for *TNFRSF1A* (A), *TGFRB1* (B), *TGFRB2* (C) and *CD44* (D).** Gene expression is presented as ln(counts per million). Yellow indicates lower expression whilst purple indicates higher expression. The left column presents moderate PCS patients and the right column presents severe PCS patients. Each point represents individual cells; n = 29.

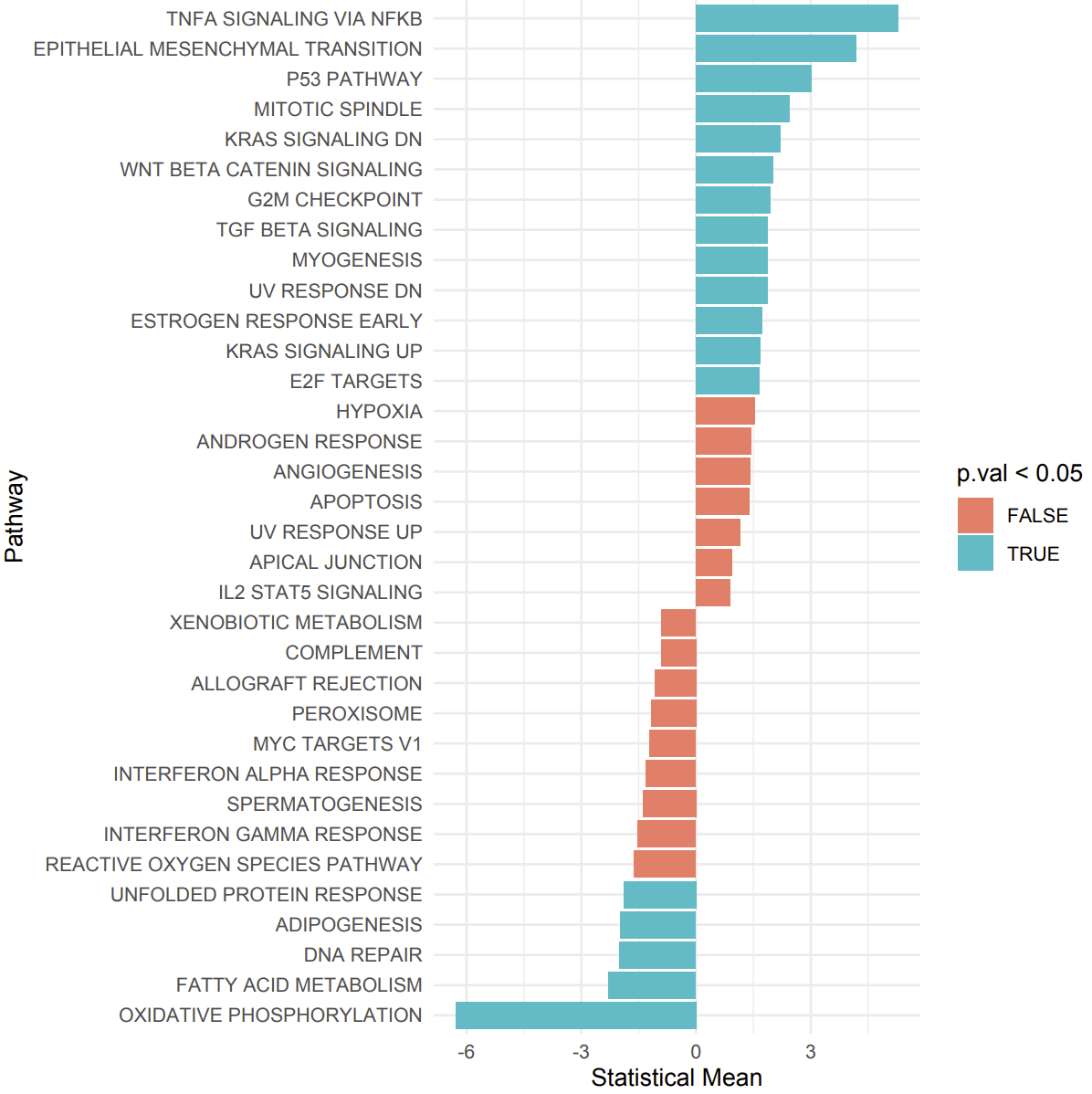

**Figure S10: GSEA of pathways enriched between moderate and severe PCS patients.** A positive statistical mean (x-axis) indicates enrichment in severe PCS while a negative statistical mean indicates enrichment in moderate PCS. Statistical significance was determined at p<0.05 indicated in blue, whilst non-significant pathways (p>0.05) are indicated in red; n = 29.

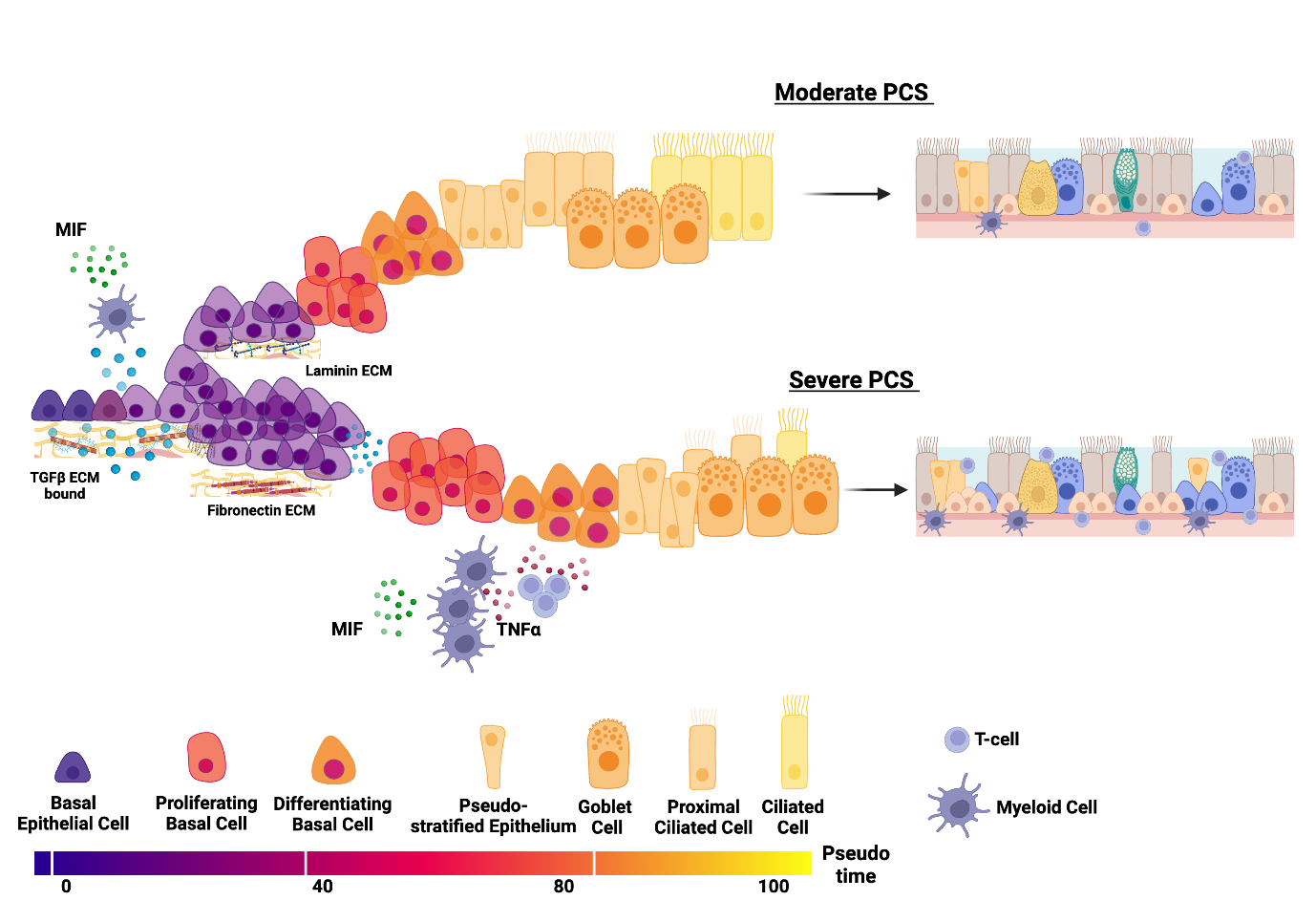

**Figure S11: Graphical summary of aberrant molecular and cellular mechanisms in the nasal epithelium in moderate vs severe PCS.**
